# Host, diet, microbiome, and xenobiotic plasma metabolites associate with kidney cell-type transcriptional programs in CKD

**DOI:** 10.64898/2026.06.11.731733

**Authors:** Farrhin Nowshad, Meltem Su, Leah Guthrie

## Abstract

**Background:** Chronic kidney disease (CKD) is associated with widespread alterations in the circulating metabolome, including metabolites influenced by diet and the gut microbiome that may represent modifiable disease pathways. However, the extent to which metabolites from distinct origins are associated with kidney cell-type-specific injury programs remains incompletely characterized.

**Methods:** We analyzed plasma metabolomics data from 240 Kidney Precision Medicine Project (KPMP) participants (207 CKD, 33 healthy reference) and validated findings in an independent cohort (n=883). We applied unsupervised clustering to identify metabotypes and elastic net classification to generate a continuous metabotype score. We classified metabolites by origin (host-derived, diet-derived, microbiome-dependent, xenobiotic) and trained random forest models to predict filtration markers and structural injury. In 66 participants with matched kidney single-cell RNA sequencing, we regressed podocyte and proximal tubule pseudobulk expression against plasma metabolites and computed hub scores quantifying transcriptional connectivity.

**Results:** Two reproducible metabotypes separated CKD from healthy states (elastic net CV AUC=0.984 discovery, 0.997 validation), with the metabotype score strongly correlating with eGFR (ρ=−0.79). Although they represented a minor subset of the circulating metabolome, diet-and microbiome-dependent metabolites such as myo-inositol, guaiacol sulfate, and indole derivatives were among the strongest metabotype-defining features. Metabolites improved prediction of filtration markers beyond clinical covariates, most notably for BUN (ΔR²=+0.29), driven by urea, guanidinosuccinate, and the microbiome-derived arabitol/xylitol. Plasma metabolites were associated with interstitial fibrosis (415 metabolites) and tubular atrophy (505 metabolites) and distinguished opposing proximal tubule transcriptional programs. Across kidney cell types, environmental xenobiotics (PFHxS, halogenated benzoic acids) activated aryl hydrocarbon receptor, inflammatory, and fibrotic programs in podocytes, while endogenous metabolites engaged inflammatory and oxidative stress programs in proximal tubule.

**Conclusion:** Plasma metabotypes capture metabolic heterogeneity beyond eGFR, with metabolites of distinct origins associated with cell-type-specific kidney injury programs, supporting plasma metabolomics as a noninvasive approach for molecular patient stratification in CKD.

**Key Points:**

- Unsupervised clustering defined reproducible plasma metabotypes spanning host-, diet-, microbiome-, and xenobiotic-derived metabolites that separated CKD from healthy reference states.
- Diet- and microbiome-dependent metabolites were among the strongest metabotype-defining features and showed the strongest associations with podocyte and proximal tubule gene expression.
- Metabolites of distinct origins associated with different cell-type transcriptional programs in podocytes and proximal tubule.

## Introduction

Chronic kidney disease affects more than 800 million people worldwide and is associated with heightened risks of cardiovascular disease, metabolic complications, and premature mortality^1–4^. Dietary patterns are an important and potentially modifiable determinant of CKD risk and treatment^5–7^, in part through their effects on the composition of the circulating metabolome. The systemic metabolome comprises small molecules originating from endogenous host metabolism, dietary exposures, and gut microbial metabolism, and serves as an integrated readout of environmental, microbial, and host physiological processes. Numerous circulating metabolites have been associated with CKD progression and cardiovascular disease risk^8–10^, suggesting that the metabolome captures biologically relevant pathways contributing to disease pathogenesis. Because diet and the gut microbiome are major determinants of many circulating metabolites^11^ and can be modified through dietary, microbial, or pharmacologic interventions, understanding how these metabolites relate to kidney injury may reveal mechanistically informed strategies to prevent or slow CKD progression.

Declining kidney function profoundly alters the circulating metabolome through impaired filtration, secretion, metabolism, and excretion, resulting in the accumulation of uremic solutes and widespread remodeling of systemic metabolic profiles^12,13^. Large-scale metabolomic studies have identified metabolites associated with eGFR decline, incident CKD, and progression to kidney failure ^13–15^. For example, Rhee et al. identified metabolites predictive of incident CKD^16^, while Sekula et al. identified metabolomic markers associated with kidney function decline and kidney failure^17^. However, most studies have focused on associations between circulating metabolites and clinical outcomes, providing limited insight into how circulating metabolites relate to cell-type-specific transcriptional programs within the kidney.

Single-cell RNA sequencing has transformed our understanding of kidney disease by revealing injury-associated transcriptional programs in distinct cell populations^18^. Studies from the Kidney Precision Medicine Project (KPMP) have identified proximal tubule states spanning healthy, adaptive, and maladaptive phenotypes and linked persistent injury programs to CKD progression^19,20^. Similarly, podocyte analyses have revealed transcriptional signatures associated with cytoskeletal dysfunction and stress responses, including activation of aryl hydrocarbon receptor signaling by uremic toxins^21–23^. Whether circulating metabolites are associated with specific injury programs in these cell populations remains incompletely understood.

Here, we leverage the KPMP cohort (n = 240) with matched plasma metabolomics, clinical phenotyping, histopathologic assessment, and kidney single-cell transcriptomic data to define the metabolic architecture of CKD. We identify reproducible plasma metabotypes through unsupervised clustering and validate them in independent cohorts (n = 883), determine the contribution of metabolites from host, exogenous, and microbiome dependent origins, to metabotype identity, evaluate associations between plasma metabolites and kidney functional and structural injury, and integrate plasma metabolomics with pseudobulk single-cell transcriptomics to identify metabolite-linked transcriptional programs in podocytes and proximal tubule cells.

## Methods

### Study Cohort and Data Sources

This study leveraged publicly available data from the Kidney Precision Medicine Project (KPMP), comprising 240 participants (207 CKD, 33 healthy reference)^24^. Plasma metabolomic data were obtained via UHPLC-MS/MS across two acquisition years and combined after log-transformation. Clinical metadata included age, sex, race, baseline eGFR, diabetes and hypertension status, and serum biomarkers (creatinine, cystatin C, BUN). Histopathological assessments of cortical interstitial fibrosis and tubular atrophy percentages were obtained from matched KPMP clinical datasets. Analyses were performed across nested subsets based on data availability: full-cohort metabolomic analyses (n = 240); metabolite-pathology and filtration marker prediction analyses (n = 83 and n = 103, respectively); and scRNA-seq integration analyses (n = 66). Characteristics of the KPMP cohort and a summary of the metabolomics data are shown in Table 1.

**Table 1.**
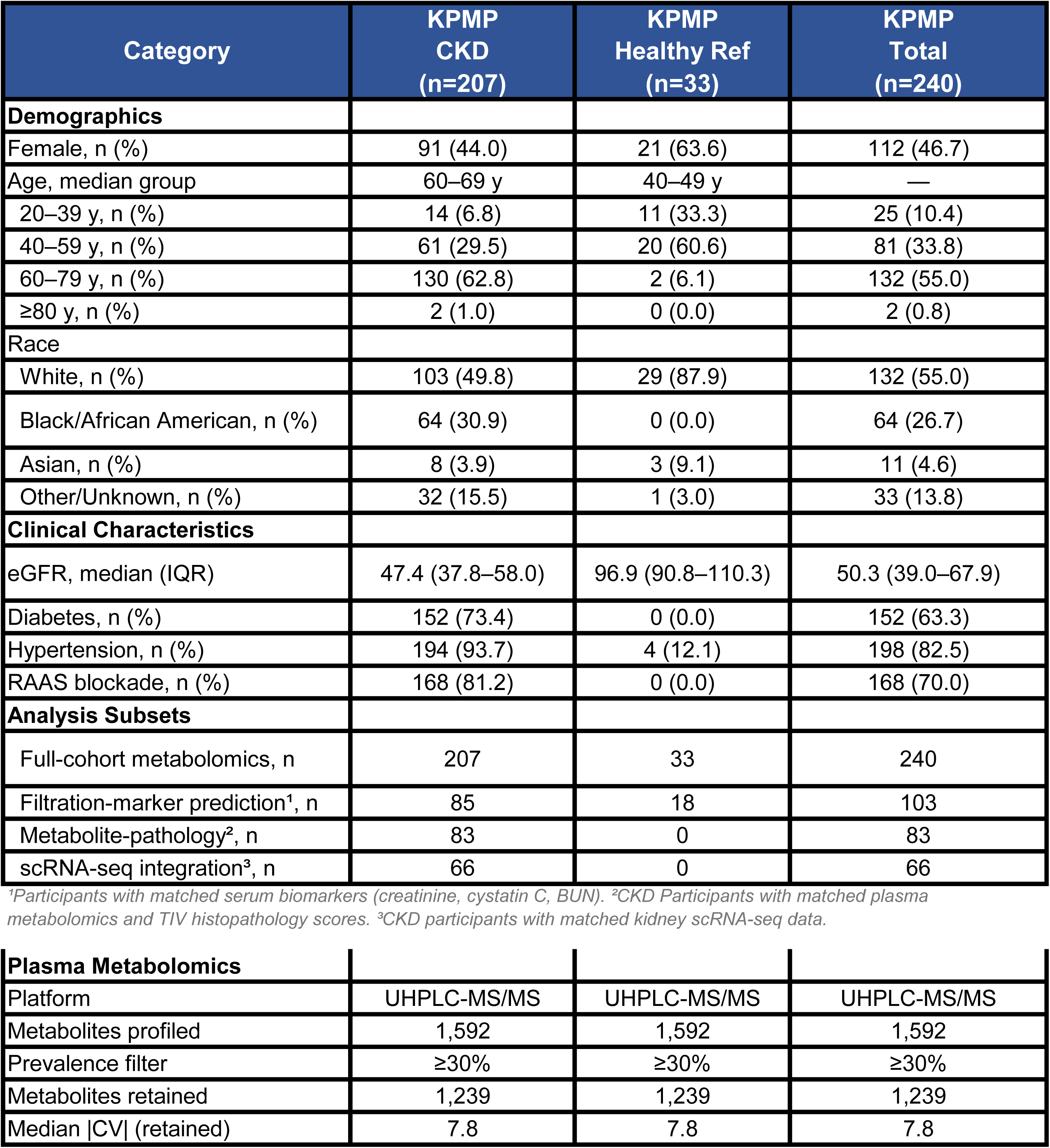
Cohort characteristics and metabolomics data summary for the KPMP discovery cohort. Demographic, clinical, and plasma metabolomics quality-control characteristics for CKD and healthy reference participants, with sample sizes for each analysis subset. CKD, chronic kidney disease; eGFR, estimated glomerular filtration rate; BUN, blood urea nitrogen; CV, coefficient of variation; IFTA, interstitial fibrosis and tubular atrophy; scRNA-seq, single-cell RNA sequencing; UHPLC-MS/MS, ultra-high-performance liquid chromatography–tandem mass spectrometry.

Findings were validated using public metabolomics data obtained using the Metabolon platform: the Modification of Diet in Renal Disease study (MDRD; n = 751 CKD) and the Assessing Long Term Outcomes in Living Kidney Donors study (ALTOLD; n = 132 healthy reference). Datasets can be found on Metabolomics Workbench (ST002818; ST002819).

### Plasma Metabolomics Quality Control

Log-transformed metabolite abundances were aggregated by median for participants with duplicate measurements. Metabolites with prevalence below 30% or zero variance were excluded, yielding 1,239 metabolites for full-cohort analyses. For the scRNA-seq matched subcohort, quality control was applied independently with a 70% prevalence threshold, yielding 1,563 metabolites. Remaining missing values were median-imputed, and the matrix was z-score standardized per metabolite. Per-metabolite quality-control metrics (prevalence, CV, mean, SD) for the KPMP dataset are provided in Supplemental Table 1.

### Unsupervised Metabolomic Clustering

Unsupervised clustering was performed using k-means (Hartigan-Wong) on the z-scored metabolite matrix. The optimal number of clusters was selected by maximizing mean silhouette width over k = 2-10 with 50 random initializations. PERMANOVA (999 permutations) tested statistical significance of cluster and enrollment group separation. PCA was performed for two-dimensional visualization with 95% confidence ellipses per cluster. Metabotype labels were ordered so that the lowest metabotype number corresponded to the group with the lowest mean eGFR. The same clustering procedure was applied to the combined validation cohort matrix using 634 shared metabolites (Supplemental Table 2). Per-participant metabotype assignments for KPMP and validation cohort participants are shown in Supplemental Table 3.

### Elastic Net Classification and Metabotype score

Metabotype classification was performed using elastic net logistic regression (glmnet, α = 0.5) with binary metabotype labels derived from k-means clustering. Model regularization was optimized by 10-fold cross-validation using AUC as the performance metric. An outer 5-fold cross-validation procedure provided unbiased performance estimates. The Metabotype score was defined as the logit-transformed predicted probability of belonging to the CKD-enriched metabotype across held-out folds. Metabolites with non-zero coefficients were defined as classifier-selected features, with coefficient magnitude and direction reflecting strength and direction of association with the CKD-enriched metabotype.

### Metabolite Origin Classification

Metabolites were classified as host-derived, diet-derived, microbiome-dependent, or xenobiotic based on prior metabolomic studies integrating bacterial culture data, human plasma profiling, and curated food metabolite databases. Metabolites with mixed or uncertain origins were assigned based on dominant evidence from biochemical annotation and pathway context^11,25^.

### Random Forest Models

Random forest regression models (ntree = 1,000) were trained to predict serum creatinine, cystatin C, and BUN from plasma metabolite profiles (n = 103). Five-fold cross-validated R² was computed for baseline covariate models (age, sex, eGFR), metabolite-only models, and combined models. Permutation importance (percent increase in MSE) identified the top 20 metabolites driving BUN prediction. Permutation importance for all plasma metabolites in the random forest model predicting serum BUN are listed in Supplemental Table 4). Separate random forest models were trained to predict interstitial fibrosis and tubular atrophy percentages (n = 83), with the same cross-validation framework applied to quantify metabolite added predictive value beyond clinical covariates; permutation importance rankings for both outcomes are provided in Supplemental Table 5.

### Metabolite-Pathology Association Analysis

Univariate linear regressions were performed for each plasma metabolite against interstitial fibrosis and tubular atrophy percentages as continuous outcomes and shown in Supplemental Table 6. P-values were corrected using the Benjamini-Hochberg FDR method; metabolites with FDR < 0.05 were considered significant.

### Single-Cell RNA-seq Pseudobulk Construction

For population-level inference across individuals in single-cell data, pseudobulk aggregation followed by bulk-style differential testing enables each participant’s cell-type-specific transcriptome to be summarized as a single expression vector that can be directly regressed against their matched plasma metabolome using established bulk RNA-seq models, while preserving valid false-discovery control across participants^26,27^. Participant-level pseudobulk expression profiles were constructed from KPMP kidney biopsy scRNA-seq data for podocytes (n = 66 participants) and proximal tubule S1/S2 cells (n = 99 participants) by averaging log-normalized gene expression across all cells per participant. Participant identifiers were harmonized between scRNA-seq metadata and plasma metabolomics data.

### Metabolite-Gene Association and Hub Score Analysis

For each metabolite-gene pair, covariate-adjusted linear regressions were performed: gene expression ∼ metabolite abundance + age + sex + eGFR, separately for podocytes and proximal tubule cells. Associations were considered significant at FDR < 0.05. Hub scores were calculated as the total number of significant gene associations per metabolite across both cell types. Genes were mapped to curated co-expression modules representing key kidney processes including inflammation, fibrosis/ECM, tubular injury, oxidative stress, podocyte identity, tubular identity, and AhR response, derived from published kidney transcriptomic literature. The dominant module per metabolite was defined as the module with the highest proportion of associated genes. Top 100 metabolites ranked by total hub score across podocyte and proximal tubule cells are shown in Supplemental Table 7.

### Metabotype Anchoring in the scRNA-seq Subcohort

In the scRNA-seq matched subcohort (n = 66), k-means clustering (k = 3) was performed on 1,563 filtered metabolites. Metabotype-defining metabolites were identified using limma linear modeling with pairwise contrasts adjusted for age, sex, and eGFR, with BH-FDR correction. Associations between metabotype and kidney function markers, histopathological measures, and transcriptional module scores were evaluated using linear regression with M1 as the reference group, adjusted for age and sex.

### Statistical Analysis

All analyses were performed in R (v4.4.1). Key packages included glmnet for elastic net classification, randomForest and ranger for random forest modeling, vegan for PERMANOVA, limma for differential metabolite analysis, Seurat for single-cell processing and module scoring, and pheatmap and ggplot2 for visualization. Fixed random seeds were used throughout to ensure reproducibility. All analysis scripts are available at https://github.com/theGuthrieLab/ckd-plasma-metabotypes.

## Results

### Cohort characteristics and study design

To investigate host-, diet-, and microbiome-dependent metabolite contributions to CKD metabolic signatures, we analyzed plasma metabolomics data from the Kidney Precision Medicine Project (KPMP) as the discovery cohort (n = 240; 207 CKD, 33 healthy references; Figure 1A; Table 1). Findings were validated in two independent publicly available metabolomics datasets: the Modification of Diet in Renal Disease Study (MDRD; n = 751 CKD) and the Assessing Long Term Outcomes in Living Kidney Donors study (ALTOLD; n = 132 healthy references). All metabolomics data was generated on the Metabolon platform and a total of 634 metabolites shared across cohorts were used for validation analyses.

**Figure 1.**
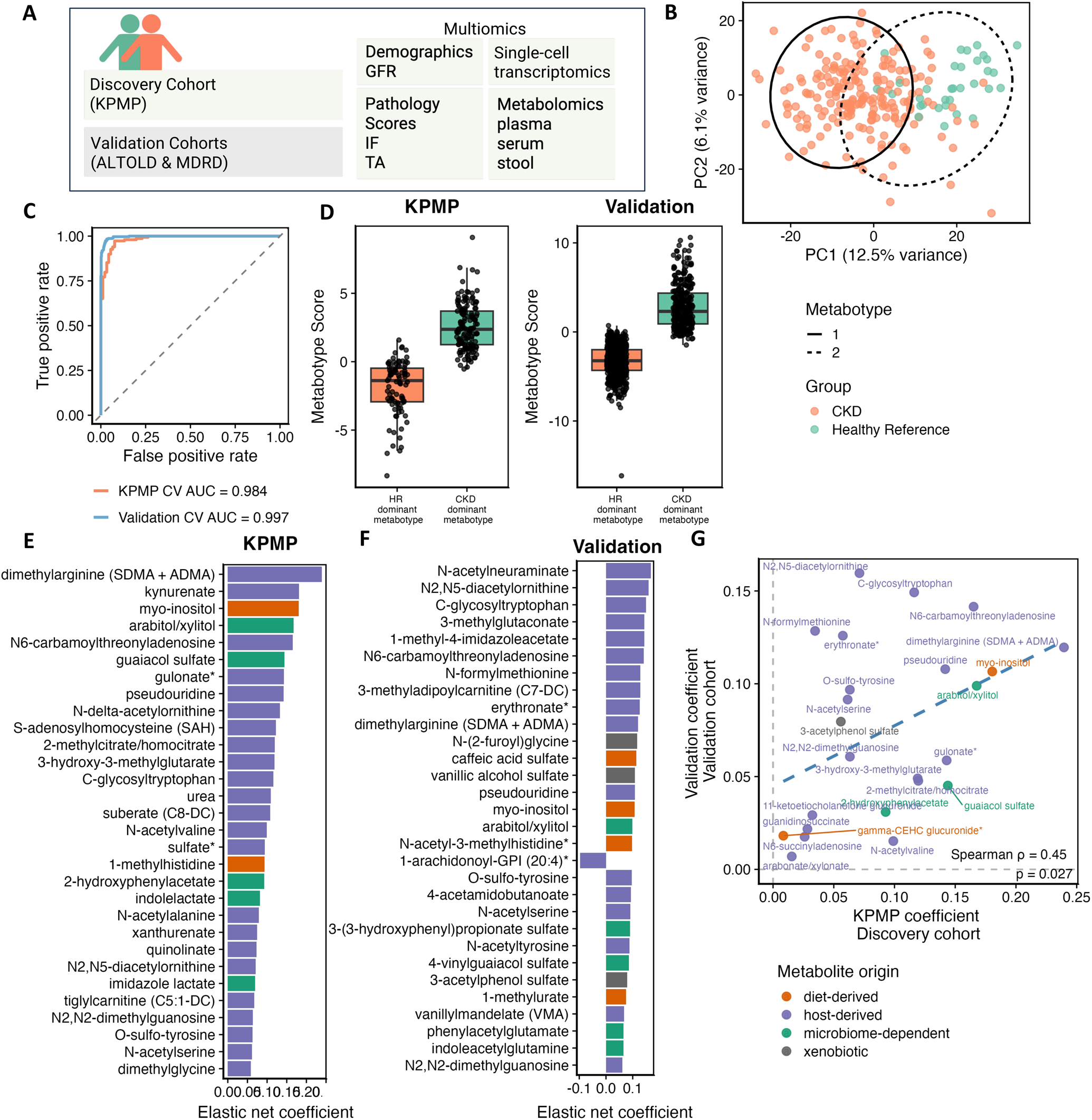
Unsupervised plasma metabolomic clustering reveals reproducible host-, diet-, and microbiome-dependent signatures distinguishing CKD from healthy individuals. (A) Study schematic showing discovery (KPMP; n = 240, 207 CKD, 33 healthy reference) and validation (ALTOLD and MDRD; n = 883, 751 CKD, 132 healthy reference) cohort composition and data types used to identify CKD metabotypes and omics signatures. (B) PCA of the KPMP plasma metabolome colored by enrollment group, with 95% confidence ellipses per metabotype (k = 2, k-means Hartigan-Wong). PERMANOVA confirms significant separation by metabotype (R² = 0.082, p = 0.001) and enrollment group (R² = 0.051, p = 0.001). (C) ROC curves for elastic net classifiers trained independently in discovery and validation cohorts to predict metabotype from plasma metabolomic profiles (CV AUC = 0.984 discovery, 0.997 validation). (D) Metabotype score distributions in discovery and validation cohorts. Metabotype score (logit-transformed cross-validated elastic net predictions) provides a continuous axis distinguishing CKD-dominant from healthy reference metabotypes. (E–F) Top elastic net model coefficients in the discovery (E) and validation (F) cohorts; bar direction indicates association with CKD-dominant versus healthy reference metabotype; color indicates metabolite origin (host-derived, diet-derived, microbiome-dependent, or xenobiotic). (G) Spearman correlation of elastic net coefficients for metabolites shared between cohorts (ρ = 0.45, p = 0.027). Concordant coefficients indicate metabolites consistently defining metabotype identity; discordant signals reflect cohort-specific effects.

### Plasma metabolomic clustering identifies reproducible CKD and healthy reference metabotypes

Plasma metabolites were clustered using an unsupervised k-means approach applied to 1, 239 quality-filtered metabolites in the KPMP dataset, resulting in two distinct metabotypes. Individuals with CKD were predominantly enriched in metabotype M1, while healthy reference participants were enriched in metabotype M2, with significant separation by both metabotype assignment (PERMANOVA R² = 0.082, p = 0.001) and disease status (R² = 0.051, p = 0.001; Figure 1B). We derived a continuous metabotype score representing each participant’s predicted probability of belonging to the CKD-enriched metabotype, based on cross-validated elastic net predictions trained on metabotype cluster assignments (Figure 1C). In both discovery and validation cohorts, healthy reference-enriched metabotypes corresponded to lower metabotype scores, demonstrating the score’s directional consistency across datasets (Figure 1D).

While unsupervised plasma metabolomic clustering has been applied in CKD, the contribution of host-, diet-, xenobiotic-, and microbiome-dependent metabolites to metabotype separation has not been systematically defined. Host-derived metabolites were the predominant metabotype score contributors. Diet- and microbiome-dependent metabolites, particularly those related to inositol, indole, and polyphenol metabolism, were also represented among the top elastic net coefficients (Figure 1E–F). In the discovery cohort, myo-inositol (diet-derived), guaiacol sulfate (microbiome-dependent), and indole metabolites ranked among the strongest contributors. In the validation cohort, top contributors included N-acetylneuraminate, N2, N5-diacetylornithine, C-glycosyltryptophan, and microbiome-dependent metabolites caffeic acid sulfate, vanillic alcohol sulfate, and phenylacetylglutamine (Figure 1F). Cross-cohort comparison of elastic net coefficients for shared metabolites demonstrated significant concordance (Spearman ρ = 0.45, p = 0.027; Figure 1G), with N2,N5-diacetylornithine, C-glycosyltryptophan, N6-carbamoylthreonyladenosine, and dimethylarginine showing consistent directionality across datasets, indicating that the metabotype score captures reproducible metabolic signatures spanning host, diet, xenobiotic-, and microbiome origins.

### Metabotype score correlates with kidney function and captures disease-associated metabolite signatures

The metabotype score showed a strong inverse correlation with baseline eGFR (Spearman ρ = −0.79; Figure 2A). Univariate linear regressions identified eGFR-associated metabolites spanning amino acid derivatives, carbohydrate intermediates, modified nucleosides, and nitrogen metabolism products, with host-derived metabolites predominating and microbiome-dependent and diet-derived metabolites also represented among the top associations (Figure 2B). Chemical class enrichment analysis revealed purine nucleotides, diazines, alkyl halides, and organooxygen compounds enriched in the CKD-dominant metabotype, while phenylpropanoic acids, coumarins, and lactams were depleted (Figure 2C).

**Figure 2.**
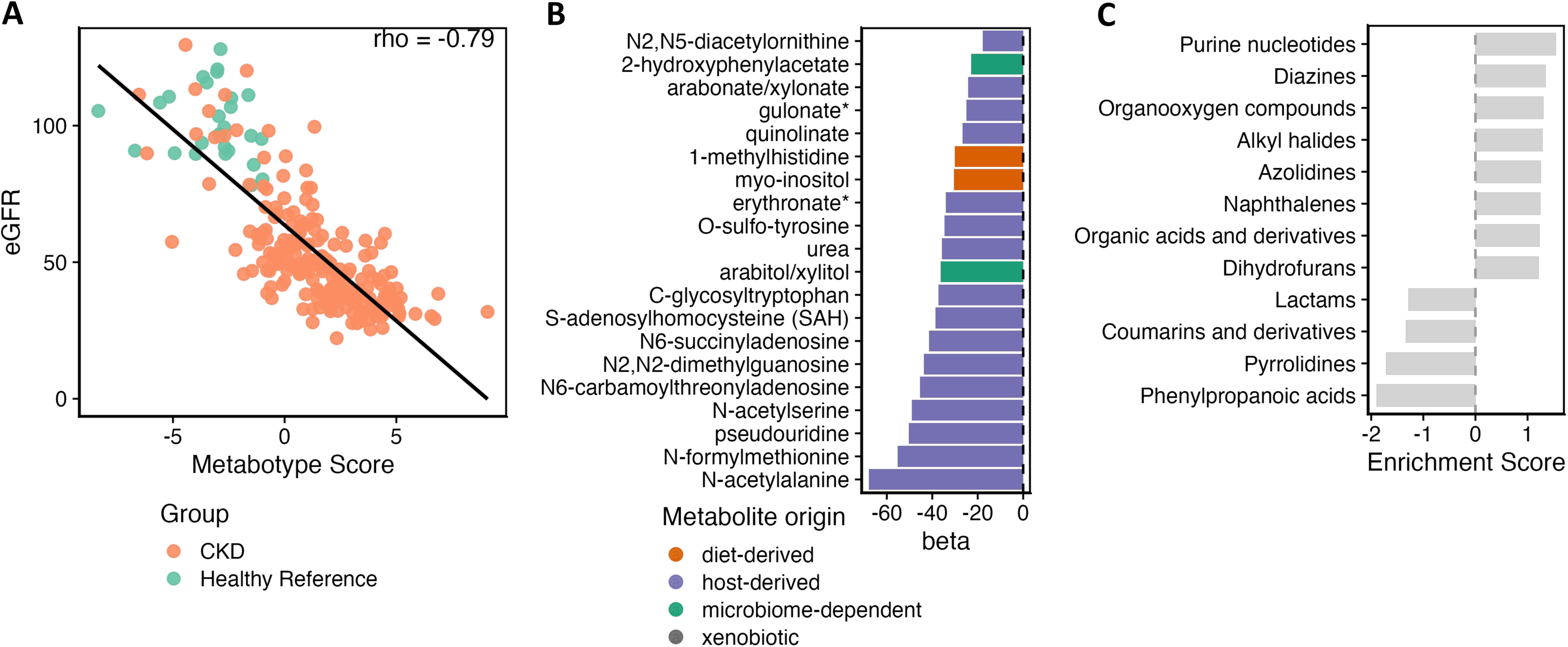
Metabotype score inversely correlates with eGFR and captures uremic metabolite accumulation and polyphenol depletion in CKD. (A) Spearman correlation between metabotype score and eGFR across CKD patients in the discovery cohort (n = 240; ρ = −0.79). (B) Bar plot of β coefficients from univariate linear regressions of eGFR on individual metabolite abundance among CKD patients; color indicates metabolite origin (C) Chemical class enrichment of metabolites associated with eGFR. Positive enrichment scores indicate classes enriched in the CKD-dominant metabotype; negative scores indicate classes enriched in the healthy reference metabotype.

### Plasma metabolites predict kidney filtration markers with highest accuracy for BUN

Building on the inverse correlation between metabotype score and eGFR, we next tested whether plasma metabolite profiles could improve prediction of kidney filtration markers. Serum creatinine, cystatin C, and BUN are complementary but each subject to well-recognized limitations; we trained random forest regression models to predict each in KPMP participants with matched serum biomarkers (n = 103).

Metabolite-only models predicted all three markers with cross-validated R² values ranging from 0.66 to 0.85 (Figure 3A). Adding metabolites to clinical covariates (age, sex, eGFR) yielded the greatest improvement for BUN (CV R² from 0.36 to 0.65; ΔR² = +0.29) and cystatin C (CV R² from 0.51 to 0.78; ΔR² = +0.27), while creatinine showed minimal gain, reflecting its role in eGFR derivation (Figure 3B). Permutation importance identified BUN prediction as driven predominantly by nitrogen-containing host-derived metabolites including urea, sulfate, N-carbamoylvaline, and guanidinosuccinate. Microbiome-dependent metabolites, including arabitol/xylitol and 3-hydroxyphenylacetylglutamine, were also among the top-ranked predictors (Figure 3C).

**Figure 3.**
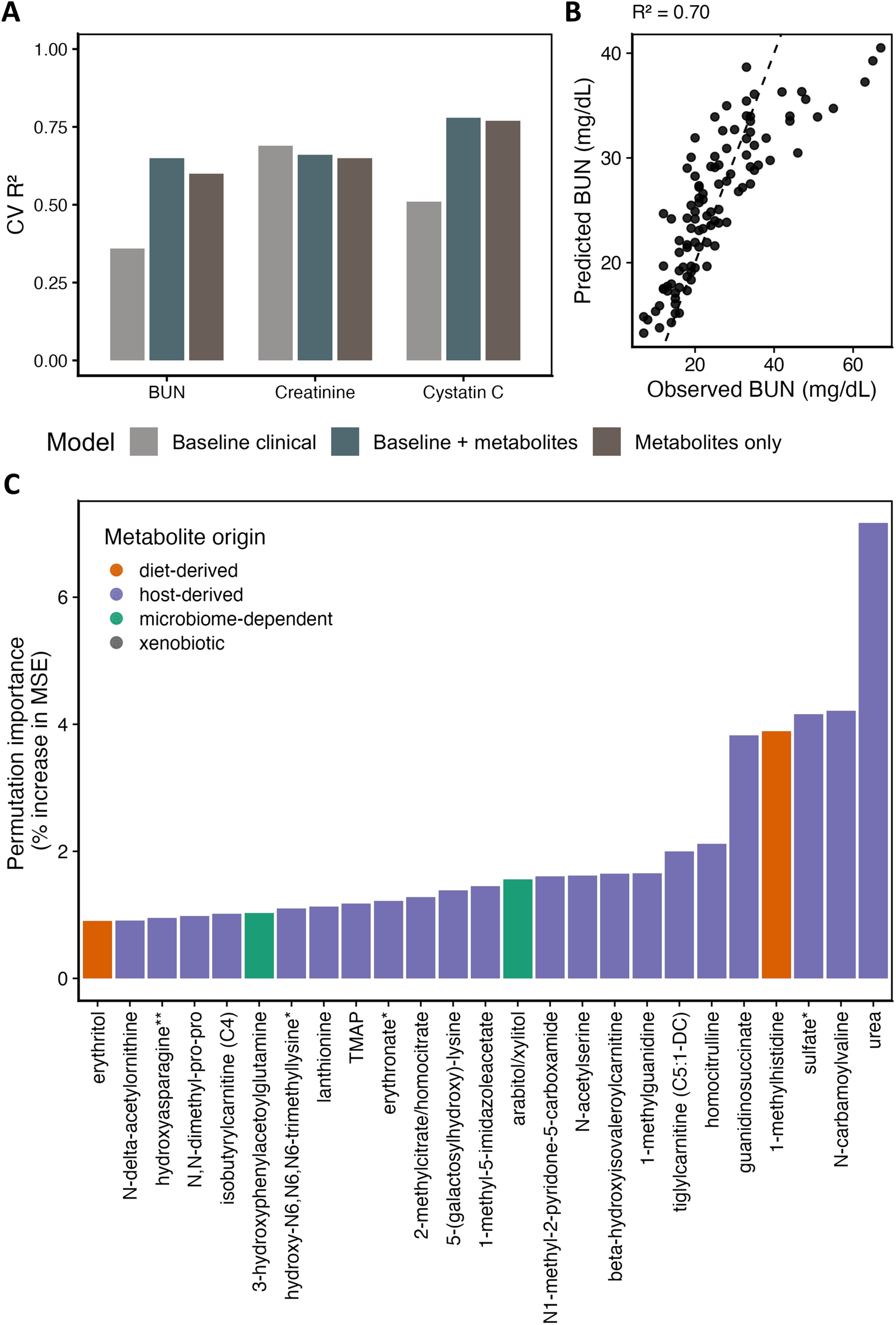
Plasma metabolite profiles predict kidney filtration markers, with highest cross-validated accuracy for BUN. (A) Five-fold cross-validated R² values for clinical baseline models (age, sex, eGFR), metabolite-only models, and combined models predicting BUN, creatinine, and cystatin C. (B) Random Forest regression model performance for predicting BUN from plasma metabolite profiles in held-out samples (R² = 0.70, RMSE = 2.79). (C) Top 20 plasma metabolites ranked by permutation importance (% increase in MSE) for BUN prediction; color indicates metabolite origin.

### Plasma metabolites associate with structural kidney injury and transcriptional connectivity hubs across kidney cell types

To investigate associations between circulating plasma metabolites and histopathological measures of structural kidney injury, we performed univariate linear regressions of plasma metabolites against cortical interstitial fibrosis and tubular atrophy percentages in KPMP participants with matched pathology scores (n = 83; Figure 4A–B). A total of 415 and 505 metabolites were significantly associated with interstitial fibrosis and tubular atrophy respectively (FDR < 0.05), with substantial overlap between the two injury measures. IFTA-associated metabolites spanned all metabolite origins, with host-derived metabolites predominating and microbiome-dependent and diet-derived metabolites represented among the most significant associations.

**Figure 4.**
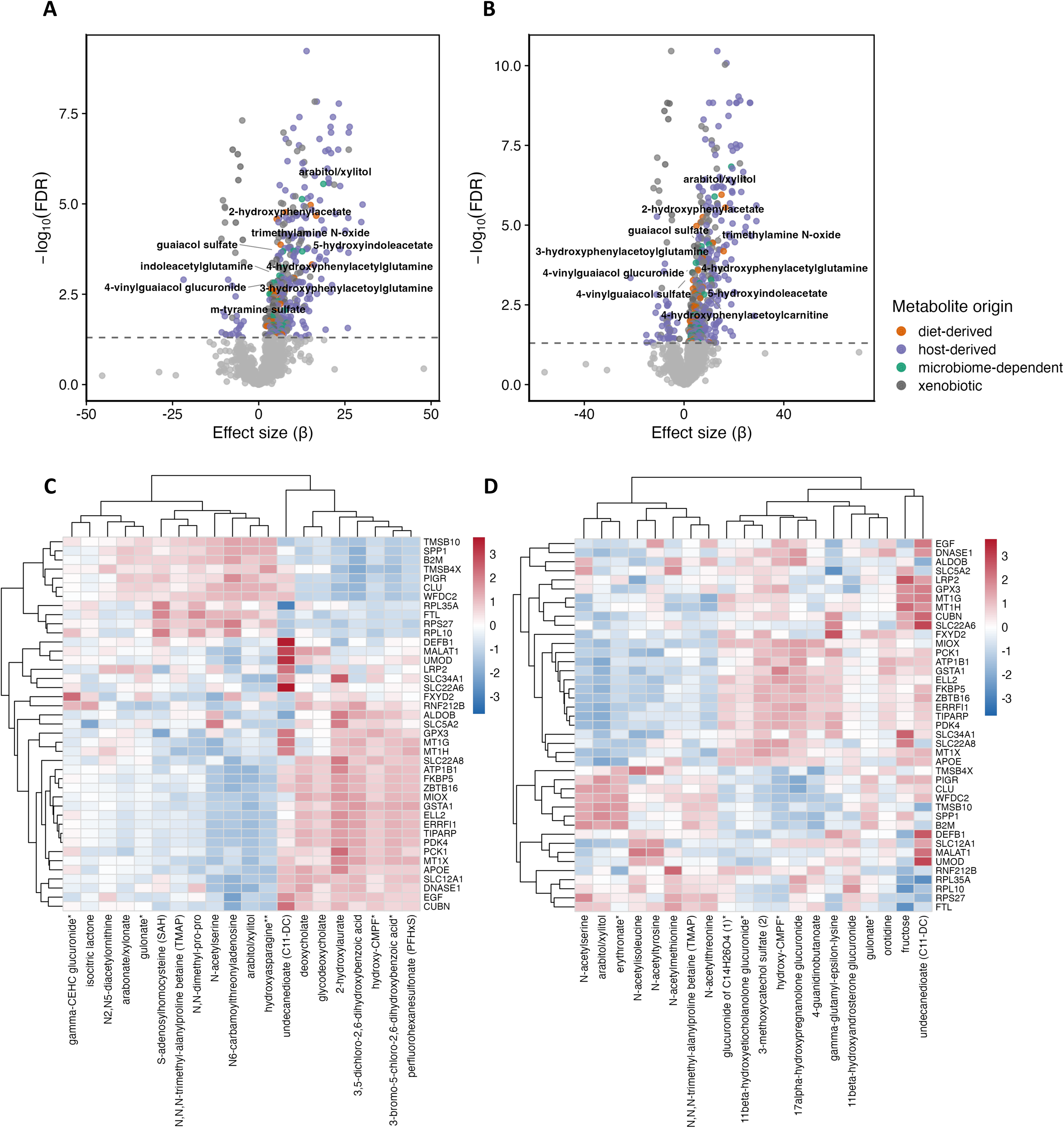
Plasma metabolites associate with interstitial fibrosis and tubular atrophy severity and link to proximal tubule transcriptional programs. (A) Volcano plot of univariate metabolite associations with interstitial fibrosis percentage (n = 83). Significant metabolites (FDR < 0.05) are colored by origin: host-derived, diet-derived, microbiome-dependent or xenobiotic; non-significant metabolites are shown in light gray. (B) Volcano plot of univariate metabolite associations with tubular atrophy percentage, colored as in (A). (C) Heatmap of row-scaled β coefficients from participant-level regressions linking the top 20 random forest-prioritized fibrosis-associated metabolites to proximal tubule (PT S1/S2) pseudobulk gene expression. Metabolites and genes are grouped by hierarchical clustering (Euclidean distance, complete linkage). (D) Heatmap of row-scaled β coefficients linking the top 20 random forest-prioritized tubular atrophy-associated metabolites to PT pseudobulk gene expression, as in (C).

To link these metabolite associations to cell-type-specific transcriptional changes, we regressed proximal tubule pseudobulk gene expression against the top 20 random forest-prioritized metabolites for fibrosis and atrophy separately (Figure 4C–D). Hierarchical clustering revealed two metabolite groups with opposing transcriptional associations: uremic retention solutes and carbohydrate metabolites associated with increased PT injury gene expression and decreased PT identity gene expression, while environmental and xenobiotic metabolites displayed the reverse pattern. Atrophy-associated metabolites were enriched for N-acetylated amino acids and steroid glucuronides, with convergent effects on PT identity and injury marker expression consistent with the fibrosis pattern.

To systematically quantify transcriptional connectivity of individual metabolites across kidney cell types, we computed hub scores capturing the aggregate strength of metabolite associations with pseudobulk gene expression across six co-expression modules in podocytes and proximal tubule cells (FDR < 0.05; Figure 5A). Host-derived metabolites comprised the largest fraction of significantly associated metabolites, followed by xenobiotic, diet-derived, and microbiome-dependent metabolites (Figure 5B).

**Figure 5.**
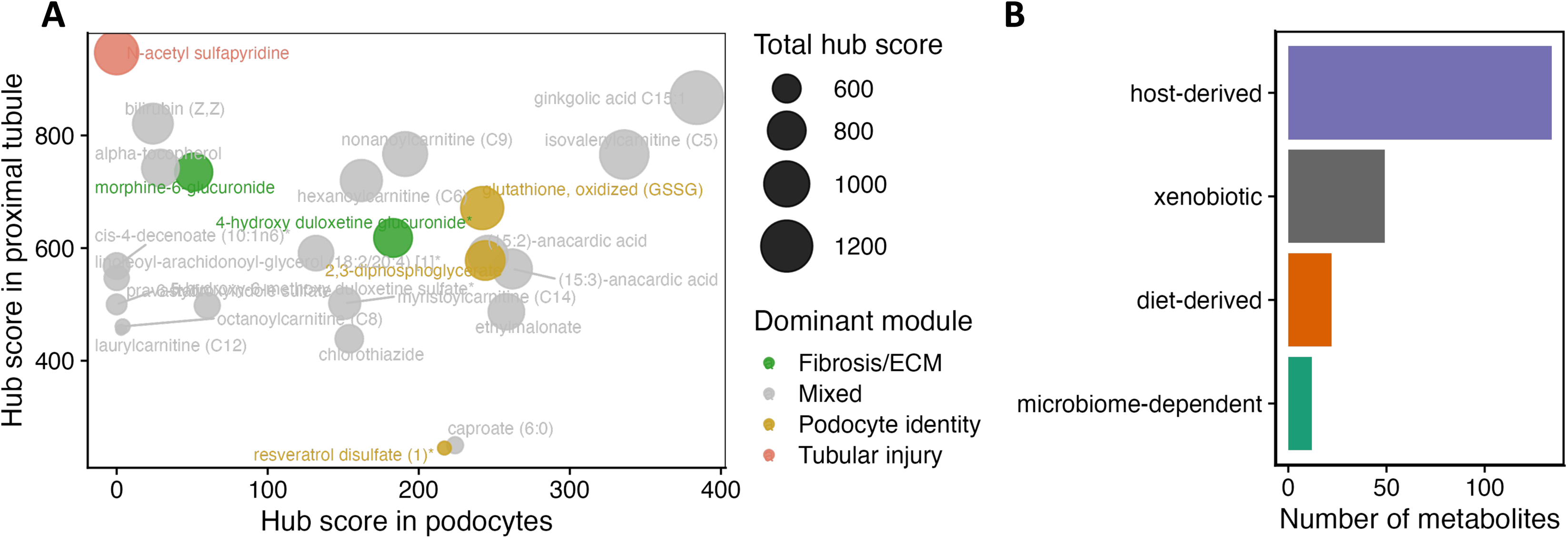
Metabolite-gene association analysis reveals cell-type-specific transcriptional connectivity hubs in podocytes and proximal tubule cells. (A) Bubble plot of metabolite hub scores derived from exhaustive metabolite–gene association analyses in podocytes (x-axis) and proximal tubule cells (y-axis) (FDR < 0.05). Each point represents a metabolite; point size reflects total hub score; color indicates dominant associated co-expression module (Fibrosis/ECM, Podocyte identity, Tubular injury, or Mixed). Metabolites positioned along both axes represent cross-compartment molecular connectors. (B) Bar plot showing the frequency of significantly associated metabolites by origin: host-derived, diet-derived, xenobiotic, and microbiome-dependent.

### CKD plasma metabotypes stratify kidney function, histologic injury, and cell-type-specific transcriptional programs

To characterize whether systemic metabolic heterogeneity within CKD maps onto distinct kidney molecular states, we performed unsupervised k-means clustering (k = 3) of plasma metabolomics data from the subset of KPMP CKD participants with matched kidney scRNA-seq data (n = 66 of 207). Three plasma metabotypes (M1–M3) were identified from 1,563 filtered metabolites, with partial separation along PC1 and PC2 (Figure 6A). Among this subset, metabotypes robustly stratified kidney function (Figure 6B): M2 exhibited markedly lower eGFR than M1 (β = −45.49, p = 7.31 × 10⁻¹¹) with significantly elevated creatinine, cystatin C, and BUN, and M3 showed comparable eGFR reduction (β = −45.86, p = 7.85 × 10⁻⁹); all associations persisted after adjustment for age and sex. In the subset with available pathology scores (n = 50), M2 and M3 showed significantly greater IFTA and tubular atrophy relative to M1 (Figure 6C). Transcriptional module scores derived from scRNA-seq pseudobulk data showed directionally consistent differences across metabotypes, with increased inflammation and decreased podocyte identity in M2, though effect sizes were modest at this sample size (Figure 6D).

**Figure 6.**
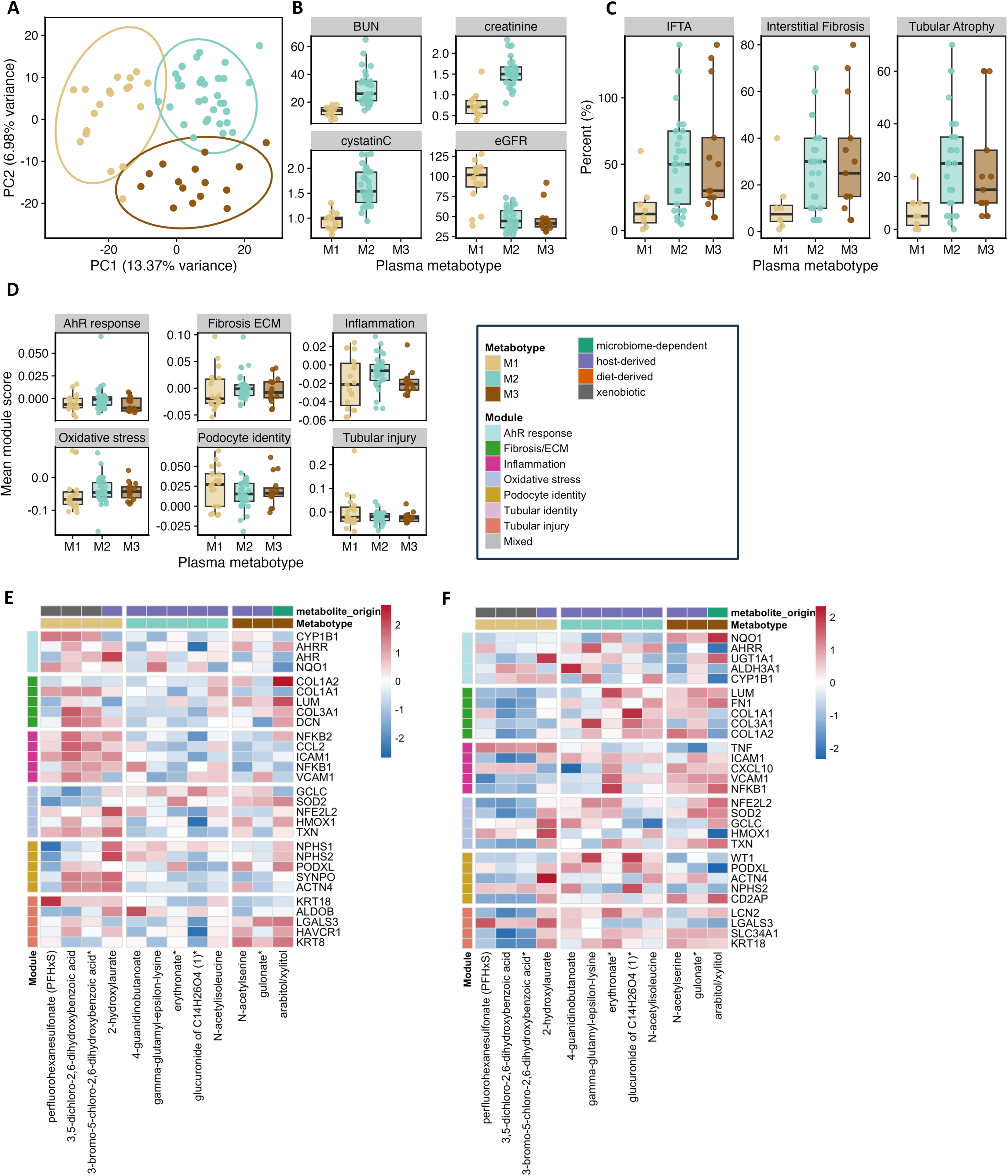
CKD plasma metabotypes stratify kidney function, structural injury, and cell-type-specific transcriptional programs. (A) PCA of scaled plasma metabolite abundances from CKD participants with matched kidney scRNA-seq data (n = 66), colored by k-means metabotype assignment (k = 3, M1–M3). (B) Kidney function markers (eGFR, creatinine, cystatin C, BUN) compared across metabotypes (M1 as reference). (C) Histologic injury measures (IFTA composite, interstitial fibrosis percentage, tubular atrophy percentage) compared across metabotypes, adjusted for age and sex. (D) Participant-level transcriptional module scores from scRNA-seq pseudobulk data compared across metabotypes for six co-expression programs. (E–F) Heatmaps of row-scaled β coefficients (adjusted for age, sex, eGFR) linking metabotype-defining metabolites to pseudobulk gene expression across six co-expression modules in podocytes (E) and proximal tubule cells (F); color in the legend indicates metabolite origin for heatmap annotations.

To identify candidate molecular pathways linking plasma metabotypes to kidney cell biology, we regressed pseudobulk gene expression against metabotype-defining metabolites across co-expression modules in podocytes and proximal tubule cells (Figure 6E–F). In podocytes, M1-associated environmental and xenobiotic metabolites showed positive associations with AhR-response and fibrosis/ECM module genes, and negative associations with podocyte identity genes, a pattern consistent with coordinated xenobiotic-associated transcriptional responses. In proximal tubule cells, M1 xenobiotics were similarly associated with AhR-response genes across both cell types; however, fibrosis, inflammation, and oxidative stress modules showed stronger associations with M2 and M3 carbohydrate and amino acid metabolites, recapitulating the tubular identity-loss pattern observed in Figure 4. Together, these exploratory findings suggest that metabolites of distinct origins may be associated with different cell-type-specific transcriptional programs: xenobiotic metabolites with AhR-mediated signaling, and endogenous uremic metabolites with inflammatory, fibrotic, and oxidative stress programs across kidney cell types.

## Discussion

This study aimed to identify plasma metabolome signatures distinguishing CKD from healthy individuals and to characterize the extent to which host-, diet-, xenobiotic-, and microbiome-dependent metabolites contribute to this separation and associate with kidney injury phenotypes. While previous studies have identified uremic toxins, altered amino acid metabolism, and gut-derived solutes as hallmarks of CKD, few have systematically resolved metabolite contributions by origin or linked plasma metabolic states to cell-type-specific transcriptional programs in the kidney. Our findings indicate that diet- and microbiome-dependent metabolites are concentrated among the strongest metabotype-defining signals despite representing a minority of the circulating metabolome, that plasma metabolites predict kidney filtration markers and histopathological injury beyond clinical covariates, and that metabolites of distinct origins are associated with different cell-type-specific transcriptional programs in this cohort, a dimension of CKD heterogeneity not captured by conventional clinical staging.

The metabotype score correlated strongly and inversely with eGFR (ρ = −0.79), yet its top-ranked metabolites span multiple dimensions of nephron handling rather than filtration alone. These include freely filtered solutes (urea, pseudouridine), protein-bound or transporter-handled solutes cleared by OAT1/OAT3-mediated secretion, and solutes normally reclaimed by proximal tubule reabsorption (myo-inositol, erythronate)^28–30^. The accumulation of secreted and reabsorbed solutes, consistent with the negative associations between uremic metabolites and proximal tubule transporter genes in our metabolite-gene analyses, is consistent with impaired tubular secretion and reabsorption beyond reduced filtration. The metabotype score therefore reflects a multi-dimensional measure of nephron function rather than a GFR surrogate, which may explain the CKD heterogeneity it captures beyond eGFR.

Linking IFTA-associated metabolites to proximal tubule pseudobulk gene expression revealed two opposing patterns. Uremic retention solutes and carbohydrate metabolites associated with increased PT injury gene expression and decreased PT identity gene expression, while xenobiotic metabolites displayed the opposite pattern. The xenobiotic-PT identity association likely reflects reverse causation: patients with preserved PT function retain the machinery to handle xenobiotic compounds, so the association marks intact tubular phenotype rather than a protective xenobiotic effect. The fibrosis and atrophy heatmaps showed convergent transcriptional consequences along the same injury-identity axis, suggesting that IFTA-associated metabolite accumulation converges on a shared dedifferentiation program regardless of whether fibrosis or atrophy is the primary lesion. The metabotype-anchored heatmaps and hub score analysis suggest that metabolites of different origins are associated with distinct cell-type-specific injury programs. In podocytes, M1-associated xenobiotic metabolites showed positive associations with AhR-response, inflammatory, and fibrosis/ECM module genes, and negative associations with podocyte identity genes^21,22,31,32^. AhR is recognized as a mediator of podocyte stress, with indoxyl sulfate-induced AhR activation triggering proinflammatory phenotypes in experimental models^33^. Our findings extend this paradigm by suggesting exogenous xenobiotics, including PFAS compounds, as candidate activators of podocyte AhR signaling in human CKD, consistent with epidemiological data linking PFAS exposure to impaired kidney function. Whether these associations reflect direct nephrotoxicity, differential PFAS clearance with declining kidney function, or environmental co-exposures warrants further investigation. The hub score analysis further identified that microbiome-dependent and diet-derived metabolites were concentrated among high-connectivity metabolites despite constituting a minority of the circulating metabolome. The depletion of polyphenol-derived metabolite classes in CKD metabotypes may reflect reduced dietary intake, impaired gut microbial polyphenol metabolism, or both, with implications for dietary interventions targeting the gut-kidney axis. These findings position diet and microbiome as priority targets for therapeutic investigation given their concentrated biological influence relative to their circulating abundance.

Several considerations help frame the interpretation of these findings and motivate future work. Because the analysis is cross-sectional, it establishes associations rather than causal relationships, and longitudinal profiling will be needed to determine whether metabotype transitions predict disease progression. The scRNA-seq–matched subcohort (n = 66) limited statistical power for transcriptional module score analyses, and these transcriptional associations should therefore be regarded as hypothesis-generating. Metabolite origin classifications may oversimplify metabolites with multiple biosynthetic sources, and univariate metabolite–pathology regressions were unadjusted for potential confounders. Transcriptional analyses focused on podocytes and proximal tubule cells, leaving thick ascending limbs, endothelial, and immune cell populations unexplored. Functional validation of the xenobiotic–AhR and endogenous metabolite–inflammatory axes using organoid or in vivo models will be important to establish causality.

In conclusion, this study shows that unsupervised plasma metabolomic clustering defines reproducible CKD metabotypes whose signatures span host, diet, microbiome, and xenobiotic origins, and links these metabotypes to distinct cell-type-specific transcriptional programs in the kidney. The metabotype score provides a continuous, noninvasive measure of CKD metabolic burden that stratifies patients along functional, structural, and molecular dimensions beyond conventional staging. The concentrated transcriptional influence of diet- and microbiome-dependent metabolites, alongside the identification of additional environmental xenobiotics as candidate activators of podocyte AhR signaling, identifies modifiable axes of CKD pathobiology that warrant further experimental and clinical investigation.

## Acknowledgements

The authors thank the Kidney Precision Medicine Project (KPMP) for providing the data used in this study. The results here are in whole or part based upon data generated by the Kidney Precision Medicine Project. Accessed January 12, 2026, https://www.kpmp.org. The Kidney Precision Medicine Project (KPMP) is supported by the National Institute of Diabetes and Digestive and Kidney Diseases (NIDDK) through the following grants: U01DK133081, U01DK133091, U01DK133092, U01DK133093, U01DK133095, U01DK133097, U01DK114866, U01DK114908, U01DK133090, U01DK133113, U01DK133766, U01DK133768, U01DK114907, U01DK114920, U01DK114923, U01DK114933, U24DK114886, UH3DK114926, UH3DK114861, UH3DK114915, and UH3DK114937. We gratefully acknowledge the essential contributions of our patient participants and the support of the American public through their tax dollars.

## Data Sharing Statement

All data used in the study are public. All analysis scripts are available at https://github.com/theGuthrieLab/ckd-plasma-metabotypes.

## Supplemental Material

**Supplemental Table 1.** Metabolite quality control for the KPMP cohort: detection prevalence, coefficient of variation, mean, and standard deviation for each plasma metabolite.

**Supplemental Table 2.** Metabolite quality control for the validation cohorts: mean, standard deviation, and coefficient of variation for each shared plasma metabolite.

**Supplemental Table 3.** Per-participant metabotype assignments for all KPMP discovery participants (n = 240) and validation cohort participants (ALTOLD, n = 132; MDRD, n = 751).

**Supplemental Table 4.** Random forest permutation importance of plasma metabolites for predicting serum blood urea nitrogen.

**Supplemental Table 5.** Univariate linear regression results for plasma metabolite associations with interstitial fibrosis and tubular atrophy.

**Supplemental Table 6.** Random forest permutation importance of plasma metabolites for predicting interstitial fibrosis and tubular atrophy.

**Supplemental Table 7.** Top 100 metabolites ranked by transcriptional hub score across podocyte and proximal tubule cells.

